# Deep-fUS: Functional ultrasound imaging of the brain using deep learning and sparse data

**DOI:** 10.1101/2020.09.29.319368

**Authors:** Tommaso Di Ianni, Raag D. Airan

## Abstract

Functional ultrasound (fUS) is a rapidly emerging modality that enables whole-brain imaging of neural activity in awake and mobile rodents. To achieve sufficient blood flow sensitivity in the brain microvasculature, fUS relies on long ultrasound data acquisitions at high frame rates, posing high demands on the sampling and processing hardware. Here we develop an end-to-end image reconstruction approach based on deep learning that significantly reduces the amount of data necessary while retaining the imaging performance. We trained a convolutional neural network to learn the power Doppler reconstruction function from sparse sequences of ultrasound data with a compression factor up to 95%, using high-quality images from *in vivo* acquisitions in rats. We tested the imaging performance in a functional neuroimaging application. We demonstrate that time series of power Doppler images can be reconstructed with sufficient accuracy to detect the small changes in cerebral blood volume (~10%) characteristic of task-evoked cortical activation, even though the network was not formally trained to reconstruct such image series. The proposed platform may facilitate the development of this neuroimaging modality in any setting where dedicated hardware is not available or in clinical scanners.

## Introduction

Functional ultrasound (fUS) is an innovative imaging modality that creates brain-wide neural activity maps at micrometer and millisecond-scale resolution by tracking temporal cerebral blood volume (CBV) changes in the brain microvasculature^1^. Similar to blood oxygen level dependent functional magnetic resonance imaging (BOLD fMRI), the detected CBV signals provide an indirect measurement of local spiking activity via neurovascular coupling^2^. However, fUS has higher spatiotemporal resolution than fMRI and uses more affordable and portable equipment, opening the possibility for functional neuroimaging performed directly at the bedside^3–6^. Preclinically, fUS enables imaging of neural activity in awake and freely behaving small animals, significantly reducing the confounding factors introduced by anesthesia/sedation or physical restraint^7,8^. Furthermore, fUS has proven useful for imaging resting state and task-evoked functional connectivity in the rat and mouse brain^2,9,10^, for visualizing vascular dynamics during spontaneous rapid-eye-movement sleep^11^, and for mapping neural activation in primates during cognition tasks and visual stimulation^12–14^. In humans, fUS has been used intraoperatively for image-monitored brain tumor removal surgeries^4,5^, and in neonates to visualize epileptic activity and to measure functional connectivity through the anterior fontanel window^3,6^.

To detect hemodynamic changes in the brain microvascular network, fUS relies on highly sensitive power Doppler sequences based on the use of plane wave emissions.

Unfocused ultrasound waves insonify the entire field of view, and the received radiofrequency (RF) data from tilted plane wave emissions are re-focused (or beamformed) and coherently compounded to increase resolution and depth of penetration. This strategy makes it possible to continuously acquire long sequences of ultrasound data at high frame rates. The obtained compound Doppler signals are then processed to filter out the strong, undesired clutter originating from the tissue, and are squared and time-integrated to create power Doppler images with pixel amplitude proportional to the CBV (**Fig. 1a**).

**Figure 1:**
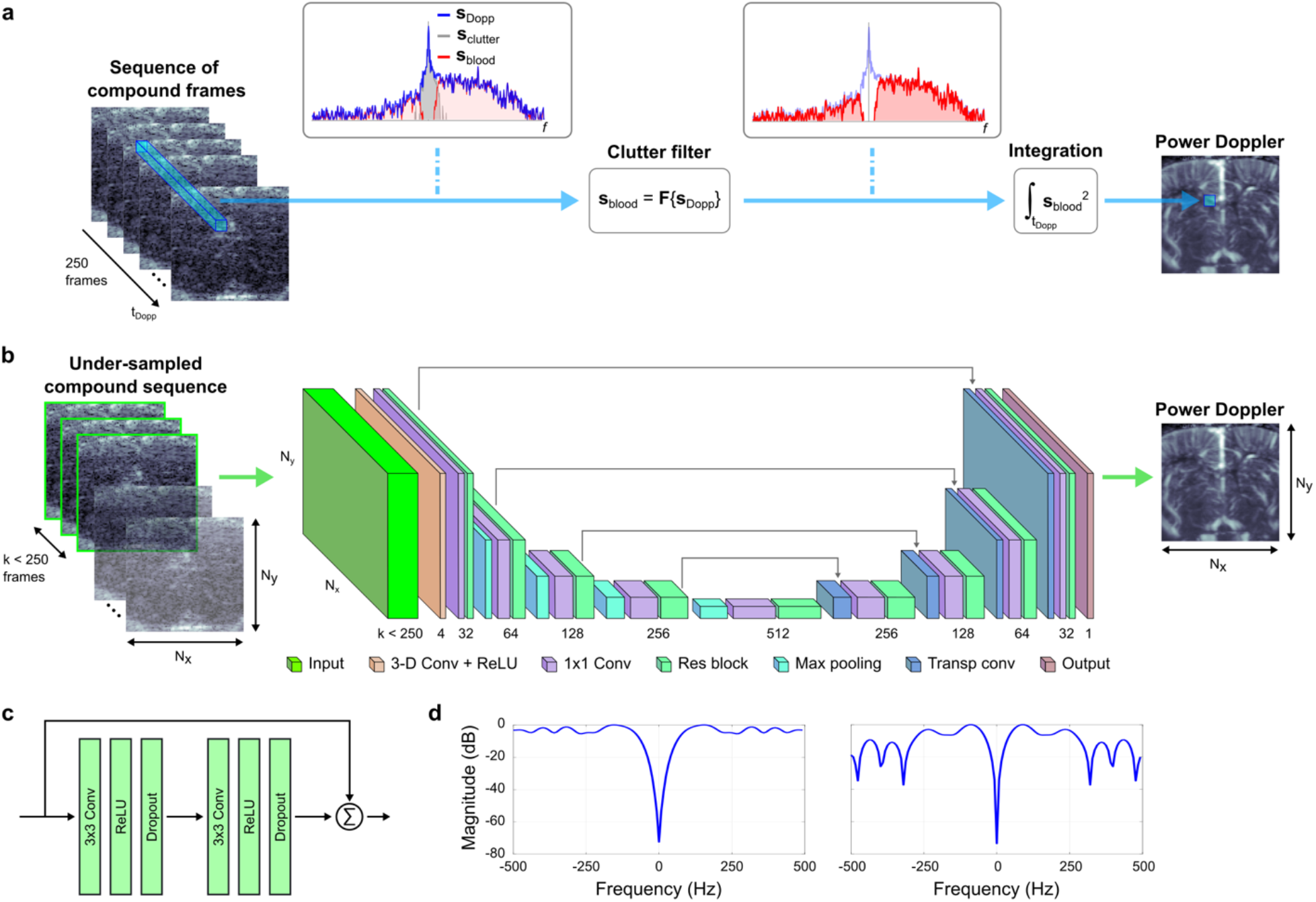
Deep convolutional neural network for the reconstruction of power Doppler images from sparse ultrasound data. **a**, State-of-the-art processing computes a power Doppler image from a sequence of 250 complex compound ultrasound frames. In a given pixel, the temporal signal **s**_Dopp_, sampled in the Doppler time t_Dopp_, is passed through a bank of filters **F** to remove the tissue clutter component **s**_clutter_. The retained blood signal **s**_blood_ is squared and time-integrated to compute the power Doppler pixel value proportional to the cerebral blood volume. **b**, The Deep-fUS architecture uses a modified U-Net network consisting of residual blocks arranged in a 5-layer encoder followed by a decoding path. An input 3-D convolutional layer extracts spatiotemporal features from the 3-D input structure. The input data is an under-sampled compound sequence created by selecting the first *k* frames of *N*_*x*_*×N*_*y*_ pixels (selected frames displayed with a green border). The network outputs the *N*_*x*_*×N*_*y*_ power Doppler image. **c**, Residual blocks are composed of two cascaded Conv/ReLU/Dropout layers and implement a shortcut connection between the input and output. **d**, Representative transfer functions of the input 3-D convolutional filters learned by the network. These were computed by performing a fast Fourier transform of the filter kernels averaged in the 3×3 spatial dimensions. The cutoff frequencies (−3 dB) for the two filters are 95 Hz (left) and 58 Hz (right).

The length of the acquisition sequence is critical to effectively discriminate the weak signals scattered by red blood cells circulating in the blood stream from the strong clutter originating in the surrounding tissue. When long observation windows are used, efficient clutter filtration can be achieved in both large and small vessels by using temporal and singular-value decomposition (SVD) filters^8,15,16^. Conversely, this filtration becomes challenging with shorter acquisitions, in particular in the smaller vessels where the blood-signal-to-clutter ratio is reduced and the low-frequency Doppler spectral components overlap with the tissue spectrum (**Supplementary Fig. 1a-b**). As a result, fUS imaging implementations use hundreds of compound frames (typically 200 to 400) to create a single power Doppler image.

The need to acquire and process large ultrasound datasets poses high demands on the hardware platform in terms of storage capacity and computational power, with data throughputs on the order of 240 MSa/image (see calculations in the Supplementary Note). These requirements make real-time fUS imaging challenging even in graphics processing unit (GPU) implementations, and such considerations are becoming increasingly relevant for volumetric fUS sequences^17,18^. Importantly, long ultrasound exposure time raises concerns about potential adverse bioeffects, even at diagnostic intensity levels^3,19^. For all these reasons, it is highly desirable to achieve state-of-the-art (SoA) fUS imaging performance with shorter ultrasound acquisitions, as this may effectively improve access to this imaging modality and expedite its clinical translation.

Here we propose an end-to-end deep learning approach to reconstruct power Doppler images from sparse compound datasets. We implemented a convolutional neural network (CNN) based on an encoder-decoder architecture (U-Net) with residual connections^20^. Variants of this model have been used for biomedical image reconstruction in applications spanning compressed sensing MRI^21^, sparse-projection photoacoustic imaging^22,23^, and sparse X-ray computed tomography^24,25^. Prior CNN applications in medical ultrasound imaging include contrast improvement^26^ and image de-speckling^27^, ultrasound contrast agent localization and tracking^28,29^, and under-sampled and adaptive beamforming^30–33^. Our network learns a reconstruction mapping between the sparse sequence of compound ultrasound data and the power Doppler output image, without requiring any prior model-based knowledge (**Fig. 1b, c**). We trained the network on high-quality power Doppler images from *in vivo* acquisitions in rats and using a custom loss function.

## Results

The received sensor RF data from 10 plane wave emissions were beamformed in a regular grid of 96×96 pixels with a spatial resolution of 100×100 µm^2^ to create compound frames. Sequences of compound frames were then processed to compute the power Doppler images (see Methods for details on the image reconstruction). The conventional processing achieves a satisfactory level of detail in coronal brain images reconstructed from 250 complex compound frames (**Fig. 2b**). The resulting SoA images were used for the CNN training and considered as a reference for evaluating the reconstruction performance. To test the power Doppler reconstruction with sub-optimal conditions, we retrospectively created sparse data sequences by selecting subsamples of *k* compound frames from each sequence, with a compression factor (CF) of 75% (*k* = 125), 80% (*k* = 100), 85% (*k* = 75), 90% (*k* = 50) and 95% (*k* = 25) (see Methods for details on the CF calculation). Power Doppler images reconstructed by the conventional processing from under-sampled data appear increasingly noisy due to the reduced blood flow sensitivity (**Fig. 2c**).

**Figure 2:**
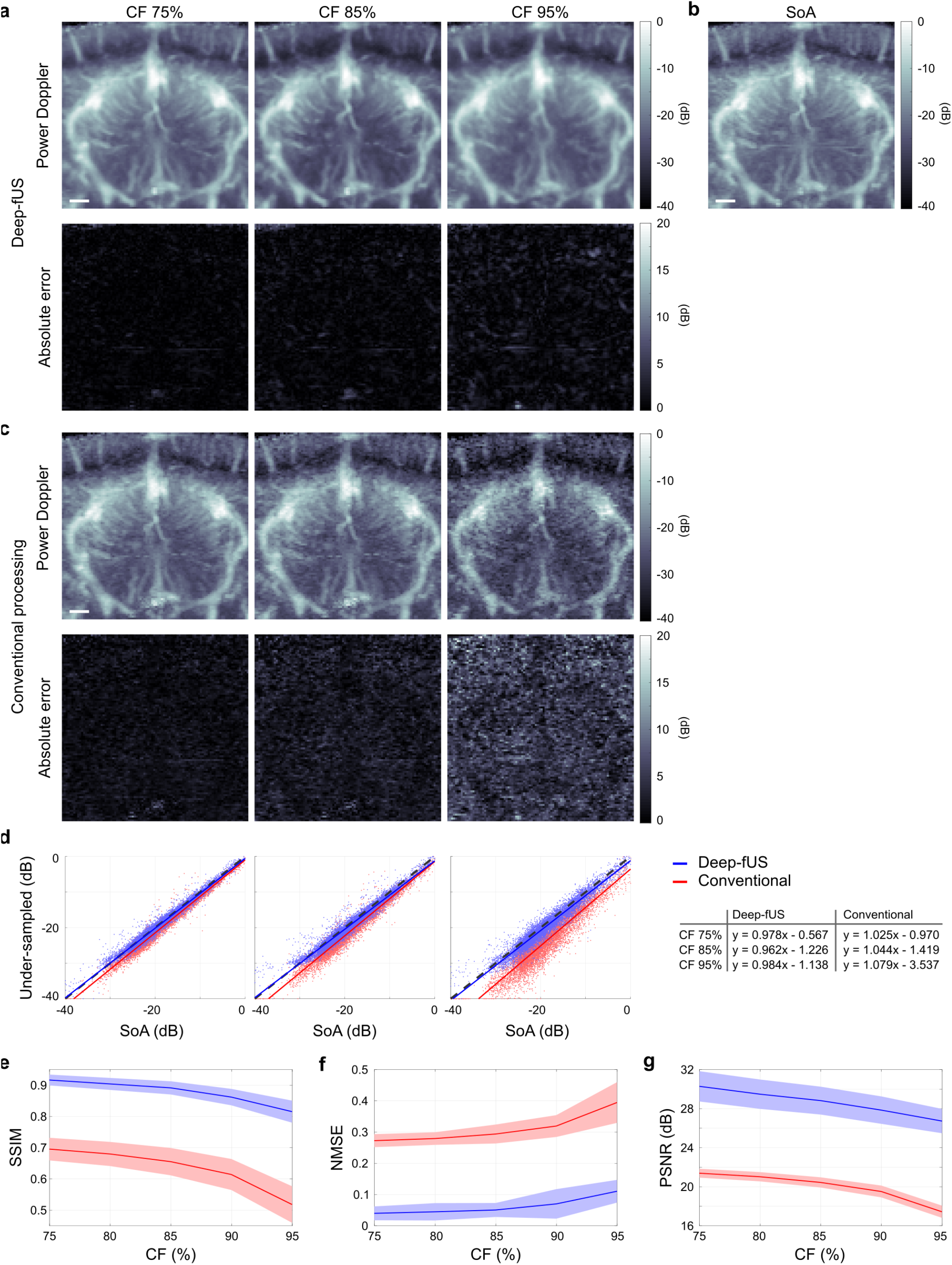
Deep-fUS produces high-quality power Doppler images from sparse sequences of compound data. **a**, Representative power Doppler image of a coronal slice of the rat brain reconstructed by Deep-fUS from under-sampled sequences with compression factor (CF) 75%, 85%, and 95% (Top) and absolute error images calculated against the state-of-the-art (SoA) image (Bottom). **b**, SoA image reconstructed by the conventional processing using 250 complex compound frames. **c**, Power Doppler images reconstructed with the conventional processing using under-sampled compound data (Top) and respective absolute error images (Bottom). **d**, Scatter plots of the power Doppler pixel amplitudes and linear regression analysis (y = b_1_ x + b_2_). **e, f, g**, Structural similarity index metric (SSIM), normalized mean squared error (NMSE), and peak signal-to-noise ratio (PSNR) of power Doppler images reconstructed by Deep-fUS (blue) and by the conventional approach (red). The quantitative metrics were calculated against the respective SoA reference images. Results are reported as mean (solid line) and standard deviation (shaded area) calculated over the test set. Scale bar in **a, b, c**: 1 mm.

The Deep-fUS network blindly solves a reconstruction problem to directly extract the power Doppler values from a sequence of compound frames. The network takes in input a sparse compound sequence and outputs the corresponding power Doppler image (**Fig. 1b**). We decided to base the processing on beamformed data instead of sensor RF data to minimize data throughput and storage (see Supplementary Note). We modified a U-Net network by (*i*) adding an input layer of 3-D convolutional filters that extract spatiotemporal features from the 3-D input data; (*ii*) using residual blocks that implement shortcut connections between the input and output features in each layer (**Fig. 1c**); and (*iii*) using dropout layers for regularization. We used a custom loss function defined as the weighted sum of the mean absolute error loss (L_MAE_) and a structural dissimilarity index metric loss (L_SSIM_) between the Deep-fUS images and the respective SoA references. We optimized relevant hyperparameters using Bayesian optimization on the dataset with CF 75%, then re-trained the network with the optimal hyperparameters and CF up to 95% (see Methods for details). Optimized hyperparameters are reported in **Supplementary Table 1**. In addition to the U-Net with residual connections (to which we refer as Res-U-Net in the Supplementary Information), we trained and optimized a network with the 3-D convolutional input layer but no residual blocks (3D-U-Net; **Supplementary Fig. c-d**) and a conventional U-Net. Furthermore, we optimized a network to perform post-processing of power Doppler images that were pre-processed using the conventional approach and sparse data (PP-U-Net). For training the reconstruction CNNs, we used a total of 740 pairs of compound data and power Doppler images from in vivo acquisitions of coronal slices of the rat brain recorded between 2.7 mm anterior and 7.04 mm posterior to bregma^34^. We used 40 additional pairs for validation and 40 pairs for testing the reconstruction performance. We based our performance analysis on the structural similarity index metric (SSIM), normalized mean squared error (NMSE), and peak signal-to-noise ratio (PSNR) between the CNN and SoA images.

The Deep-fUS reconstruction network restores the reference imaging performance and is able to reconstruct the rat brain microvasculature from sparse data with a CF up to 95% (**Fig. 2a**). Our CNN produces a considerable improvement in the under-sampled power Doppler reconstruction when compared to the conventional processing, as confirmed by the quantitative metrics in **Fig. 2e-g**. By analyzing the scatter plots, it is evident that reconstruction errors are more prominent in correspondence of the lower power Doppler values, particularly in the case of conventional processing (**Fig. 2d**). In **Supplementary Fig. 2** we display a second Deep-fUS test image, and **Supplementary Fig. 3** shows the reconstruction/post-processing results with the other trained networks. While all the trained CNNs perform significantly better than the conventional processing when sparse data are used, the Res-U-Net achieves superior reconstruction performance, with maximum SSIM of 0.92, PSNR of 30 dB, and NMSE of 0.04 (**Supplementary Table 3**). We note that the most significant improvement between the U-Net and Res-U-Net is obtained with the introduction of the 3-D convolutional input layer, which is responsible for 91% of the SSIM increase and 77% of the NMSE reduction. The mean prediction time for the Deep-fUS network is between 4.4 and 13.5 ms/image. The post-processing U-Net provides comparable imaging performance but adds a time overhead of ~210 ms/image for the pre-processing of the power Doppler images (**Supplementary Table 2**). Moreover, it is worth pointing out that this approach is similar to the conventional processing method, in that it is inherently dependent on the design of the clutter filtration stage. Interestingly, we noted that the learned convolutional filters in the input layer of the Res-U-Net implement high-pass transfer functions with strong rejection of the 0-Hz component (**Fig. 1d**). These filters seem to mimic the temporal filters used in the conventional processing but are learned directly from the data during training. Movies showing side-by-side comparisons of the conventional and Deep-fUS reconstruction approaches are available in **Supplementary Videos 1-3**.

We then sought to evaluate whether Deep-fUS provides sufficient accuracy in the reconstruction of time series of power Doppler images in a functional neuroimaging application. We imaged visual task-evoked brain activation in a rat exposed to binocular green light stimulation. We created activation maps by performing pixel-wise correlations between the power Doppler signals and the temporal stimulus pattern (**Fig. 3a**). A statistical threshold was applied to create binary masks showing only the significantly correlated pixels (P < 0.001; see Methods for details). The SoA activation map is shown for reference in **Fig. 3b**. This was created using conventionally reconstructed power Doppler images using the full compound dataset and shows significant activation in the primary and secondary visual cortices (V1M, V1B, and V2MM) bilaterally. In **Fig. 3c** we report the activation maps created using power Doppler time series reconstructed by Deep-fUS using sparse data with CF between 75% and 95%. Although the quality of the activation maps decreases with increasing data sparsity, significant visual cortex activation can be detected with a CF up to 95%. Notably, Deep-fUS with CF 95% performs better than the conventional approach with CF 75%, as confirmed by the quantitative error metric (**Fig. 3g**). With shorter data sequences, the conventional processing provides noisy CBV temporal signals that result in lower and non-significant correlations with the stimulus pattern (**Fig. 3e; Supplementary Fig. 5**). It is important to point out that although the neural network was not formally trained to reconstruct such time series of power Doppler images, it generalized well to this reconstruction task and was able to detect the small changes in relative CBV signal (~10%; **Fig. 3f**) characteristic of visual-evoked cortical activation.

**Figure 3:**
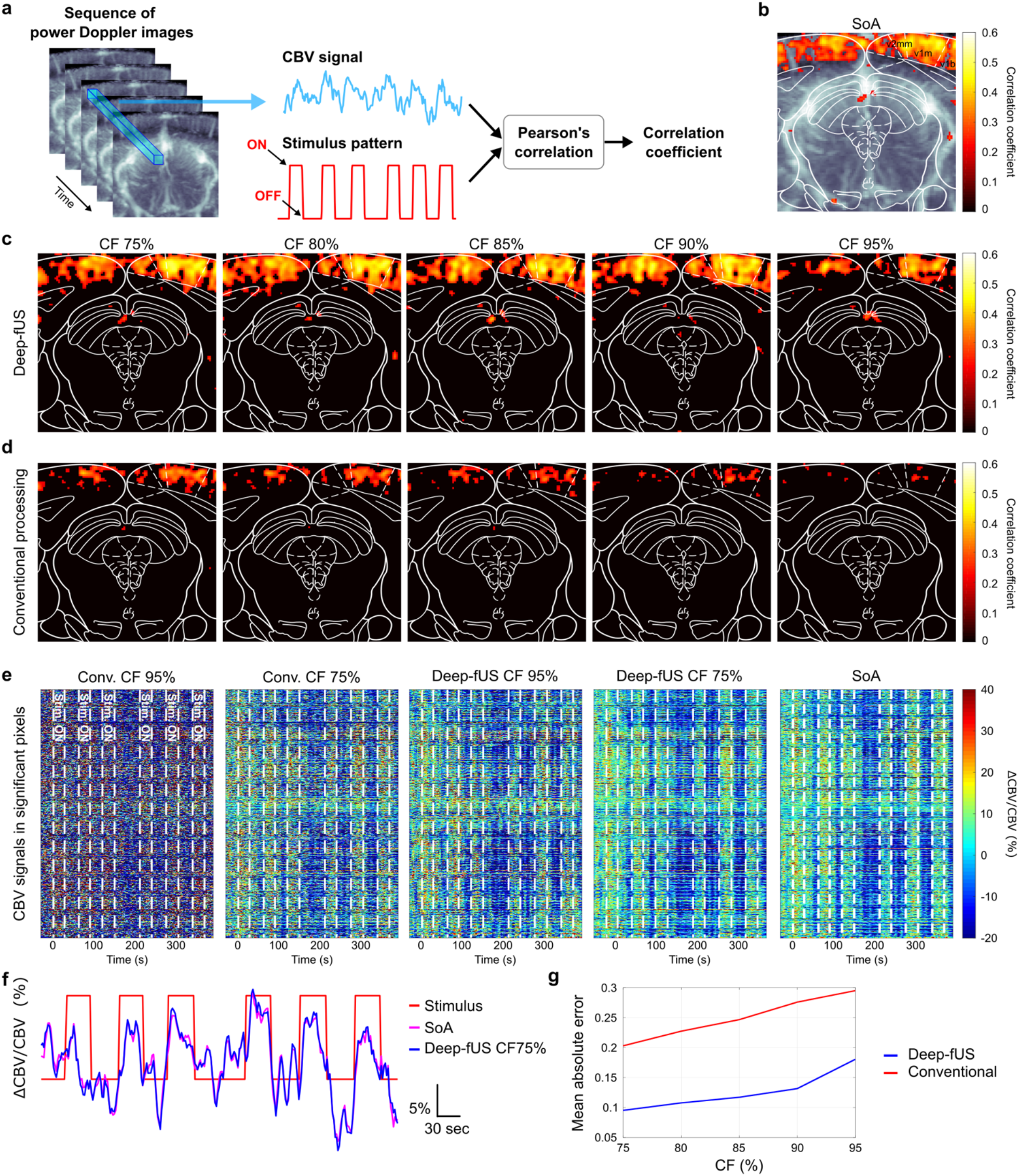
Maps of visual task-evoked brain activation computed using sparse data sequences. **a**, Time series of power Doppler images were recorded continuously during a visual stimulation task. The resulting cerebral blood volume (CBV) signals were correlated with the stimulus pattern. The stimulation consisted of 6 light stimuli, each with an ON time of 30 s, distributed in a pseudo-random fashion. **b**, State-of-the-art (SoA) activation map computed using power Doppler images reconstructed by the conventional approach using 250 complex compound frames. The activation map (heat map) is superimposed on a power Doppler image. The white contour represents the slice at bregma −7.04 mm from the Paxinos brain atlas^34^. The activation map shows significant bilateral activation of the rat primary and secondary visual cortices (V1M/V1B/V2MM). **c**, Activation maps computed using power Doppler images reconstructed by Deep-fUS with compression factor (CF) between 75% and 95%. **d**, Activation maps computed using power Doppler images reconstructed by conventional processing with CF between 75% and 95%. **e**, Relative CBV temporal signals in all the statistically significant pixels of the SoA map in **b**. The white dashed lines display the stimulus ON/OFF times. **f**, Mean relative CBV signals in the activated regions from the SoA activation map and Deep-fUS map with CF 75%. **g**, Mean absolute error between the statistically significant correlations in the SoA map and the under-sampled maps.

Next, we created temporally and spatially under-sampled sequences by retaining only a subset of compound samples in each frame, with a spatial under-sampling ratio *m* = 1/2 and *m* = 1/4 (**Supplementary Fig. 4**). We selected *k* = 50 and *k* = 100 in the two cases to equalize the CF to 95%. This approach improved the quality of the functional activation maps compared to the case with temporal under-sampling alone (**Fig. 4; Supplementary Table 4**), demonstrating that spatial sparsity may be a viable option to increase data compression while retaining the advantages of longer acquisitions. We provide a movie of the Deep-fUS power Doppler series and relative CBV variation from the visual-evoked experiment reconstructed using temporally and spatially sparse data in **Supplementary Video 4**.

**Figure 4:**
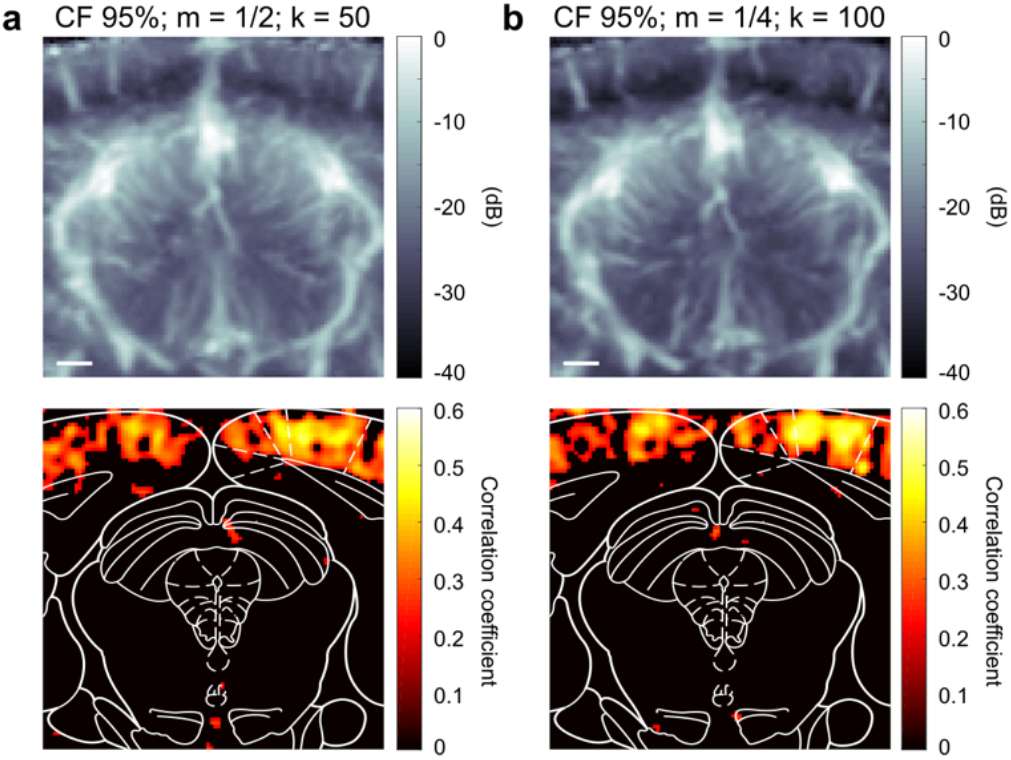
Deep-fUS reconstruction of temporally and spatially under-sampled data restores the image quality of longer acquisitions while increasing data compression. Representative power Doppler test images and activation maps computed with spatially under-sampled sequences with spatial sampling ratio m = 1/2 (**a**) and m = 1/4 (**b**). To equalize the compression factor (CF) to 95%, *k* = 50 and *k* = 100 compound frames were used in the two cases.

To investigate the effect of the acquisition length on the imaging performance in the presence of animal motion, we used Deep-fUS to reconstruct a time series of power Doppler images acquired in a lightly sedated animal. We computed the SSIM of each image in the series versus a baseline calculated as the mean of 100 images from the same acquisition, then we applied a SSIM threshold to discard the images that were significantly degraded compared to the baseline, possibly as a result of animal motion (**Fig. 5c-d, Supplementary Fig. 6**). In the case of SoA processing using data from the full compound sequence, 7.2% of the power Doppler images were discarded by the filter (**Fig. 5a**). On the other hand, shorter data sequences decrease the likelihood of animal motion during the acquisition window and reduce image scrubbing to between 2.7% (with CF 75%) and 1.2% (with CF 95%). In **Fig. 5b**, we display a representative power Doppler image that was discarded by the motion filter in the case of SoA processing but retained in all the under-sampled cases. We provide a side-by-side comparison of the SoA and Deep-fUS image scrubbing in the lightly sedated fUS experiment in **Supplementary Video 5**.

**Figure 5:**
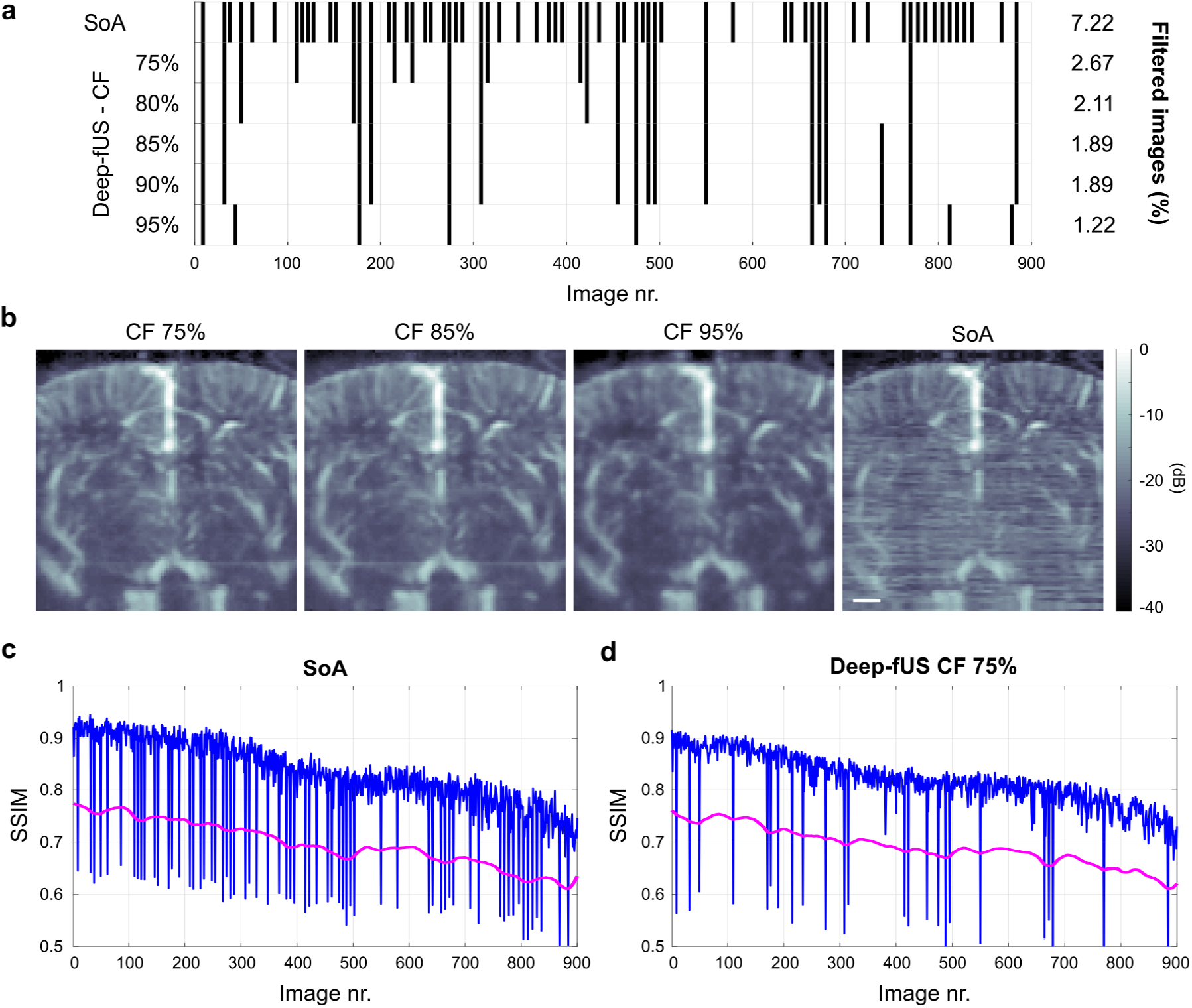
Deep-fUS with sparse sequences reduce image scrubbing due to motion artifacts in a lightly sedated functional imaging experiment. **a**, A series of 900 power Doppler images was filtered based on a structural similarity index metric (SSIM) threshold. Vertical bars in the plot show the discarded images in the series. **b**, Representative power Doppler coronal image in a case of significant degradation in the conventional reconstruction that was completely resolved in the under-sampled processing. Scale bar: 1 mm. **c, d**, SSIM for each image in the series versus a baseline calculated as the mean of 100 images from the same acquisition (blue). To filter images that were significantly degraded relative to the baseline, we used an SSIM threshold (magenta) defined as 85% of the local SSIM value.

## Discussion

Deep learning and CNNs are drawing increasing attention for the reconstruction and processing of biomedical images with sparse data^35–37^. In medical ultrasound, several strategies have been proposed to restore high image quality while reducing data sampling, transmission, and processing^30,31,33,38^. With the exception of a single preliminary study reporting deep learning of color Doppler images^39^, however, CNNs have not been applied as extensively to ultrasound imaging of blood flows. We demonstrated a deep learning method for the direct reconstruction of power Doppler images from a 3-D space of sparse compound ultrasound data (**Fig. 1**). Our proposed approach largely enhanced imaging performance compared to the conventional method with compression factors up to 95%, as clearly indicated by the presented quantitative metrics (**Fig. 2 and 3**). We expect the high quality of the *in vivo* rat brain images used during training to be a key contributing factor to the Deep-fUS reconstruction performance^22^. We demonstrate that our network is able to reconstruct time series of power Doppler images with sufficient accuracy to compute functional activation maps in a task-evoked neuroimaging application (**Fig. 3 and 4**). Although it was not formally trained on such a reconstruction task, the network generalized well and detected changes in relative CBV signals on the order of 10%. Additionally, we show that by minimizing the length of the acquisition sequence, our network allows greater robustness to motion artifacts in an experiment with a lightly sedated animal and is less sensitive to image scrubbing (**Fig. 5**).

The main advantage of using sparse sequences is the net reduction in data acquisition, storage, and processing demands. The suggested approach can facilitate the development of fUS neuroimaging in any setting where dedicated hardware is not available or in clinical scanners, making this technology more affordable and opening the way to new potential applications based on this imaging modality^3,40^. Additionally, sparse sequences may prove beneficial in experimental situations where fUS acquisitions need to be interleaved with long therapeutic ultrasound pulses, such as in the monitoring of focused ultrasound neurointerventions^41,42^. Importantly, this method significantly reduces the exposure time and lowers the risk of harmful bioeffects, making brain ultrasound neuroimaging safer^3,19^. Although in this study we retrospectively under-sampled the compound data, we clearly demonstrate that the network may considerably reduce the beamforming complexity and eliminate the need for computationally demanding filters^15,16^ (see Supplementary Note). Additionally, the network has the potential to increase the imaging frame rate and to facilitate the implementation of volumetric fUS imaging using swept linear arrays^6^. The platform and conceptual framework that we propose may be adapted to other high-frame-rate Doppler ultrasound imaging modalities, including vector flow imaging, to expedite deployment in portable ultrasound systems^43,44^. In creating sparse sequences, we decided to select only the initial portion of the original sequence instead of selecting and retaining interleaved frames to take advantage of the shorter temporal acquisition windows. This approach has the benefit of reducing the occurrence of motion artifacts and signal degradation due to data scrubbing, which are inevitable factors in mobile rodent fUS imaging experiments^8,45^ and handheld applications^5^ and are more likely to appear with longer acquisition times.

We acknowledge that variants of the U-Net have been previously applied to different biomedical imaging modalities. However, most of the literature is focused on removing artifacts from sub-optimally reconstructed images. We were specifically interested in demonstrating a data-driven reconstruction method that, once trained, requires no prior model-based knowledge of the image formation process nor requires hand-picked parameters. We decided to base our implementation on the U-Net as we hypothesized that its encoder-decoder architecture would well fit the nature of our data. A critical step in the power Doppler reconstruction process is the filtration of the strong clutter signal originating from the moving tissue. In the 3-D space formed by the image plane and Doppler time, the clutter signal is slowly varying in the temporal domain and highly correlated in the spatial domain, therefore it is crucial to account for both spatially and temporally varying features in the reconstruction process. By progressively expanding the spatial field of view in the encoder layers and with the input filters performing temporal convolutions, our network is able to efficiently extract spatiotemporal features from severely under-sampled input datasets. By using more sophisticated networks, we envision that interesting applications may be developed in the future based on the current work. Unsupervised algorithms may be designed to train variants of generative adversarial networks (for example, CycleGAN^46^) on 2-D power Doppler images for the reconstruction of volumetric fUS data acquired with sparse physical apertures. Considering the cost, complexity, and bulkiness of 3-D ultrasound systems, such advances may greatly facilitate 4-D fUS and other volumetric ultrasound imaging applications.

A main limitation of using ultrasound for brain imaging is the presence of the skull, which is highly absorbing in particular at the high imaging frequencies. This has limited clinical fUS to intraoperative applications or to scenarios with natural skull openings, such as the neonatal anterior fontanel window. The reduced data required by our proposed method could facilitate implementations of fUS with focused emissions, which may be more efficient in the presence of the skull. However, we should note that at a frequency of 15 MHz the rat skull would introduce a significant attenuation of the ultrasound energy and consequent heating of the bone. As a reference, a study found that the one-way transmission rate decreases rapidly from 91% at 0.3 MHz to 57% at 2.5 MHz, with a mean rat skull thickness of 0.7 mm^47^. In pulse-echo imaging, the ultrasound would have to cross the skull twice, and the focus will be deflected due to the non-uniform bone thickness and density. These limitations could be partly overcome by using contrast agents to enhance the ultrasound signal^48–50^.

## Methods

### Deep-fUS network

We modified a U-Net network^20^ and trained it to perform the power Doppler reconstruction task. This fully convolutional neural network is based on an encoder (contracting path) followed by a symmetrical decoder (expanding path). The contracting path progressively down-samples the input data and learns high-level features that are propagated to the following stages. The decoder uses up-sampling operators to increase the resolution of the encoder features and to consecutively restore the input resolution at the output stage. Skip connections between the encoding and decoding paths allow to retain context information, which is propagated to the symmetric up-sampling layers. In our model schematically illustrated in Fig. 1b-c, we modified the U-Net by adding an input layer of 4 3-D convolutional filters followed by rectified linear unit (ReLU) activations. The size of the filter kernels was considered as a hyperparameter and was optimized via Bayesian optimization (see “Training and hyperparameter optimization” section). This layer extracts spatiotemporal features from the 3-D input structure, and the transfer functions of the learned filters present a strong rejection of the 0-Hz component, resembling the temporal filters used in the conventional processing approach (Fig. 1d). In addition, we replaced convolutional layers with residual blocks composed of two cascaded Conv/ReLU/Dropout layers and included shortcut connections between the input and output features. Residual blocks were arranged in a 5-layer encoder followed by a 4-layer decoder path, and implement 3×3 convolutions followed by ReLU activations and a dropout layer to improve regularization. The dropout rate was considered as a hyperparameter and optimized via Bayesian optimization. We used 1×1 convolutions at the input of each layer to equalize the number of input and output features of each residual block. In the encoder path, down-sampling is performed by a 2×2 max pooling operator that halves the resolution in both the image dimensions. In the decoder path, 2×2 transposed convolutions with ReLU activations are used as up-sampling operators. The number of channels is progressively increased in the encoder path (32, 64, 128, 256, and 256 filters) and then decreased in the decoder (256, 128, 64, and 32 filters). The output layer is a single-channel 1×1 convolution block. The stride is equal to 1 in all the convolutional layers and 2 in the max pooling and transposed convolution blocks. All the convolutional kernels were initialized using the He initialization^51^. This network has a total of 9,788,421 trainable parameters (Supplementary Table 2). We refer to this network as Res-U-Net in the Supplementary Information or generally as Deep-fUS in the main text.

### 3D-U-Net, U-Net, and PP-U-Net networks

Along with the Res-U-Net described above, we trained and optimized three additional networks. The 3D-U-Net is analogous to the Res-U-Net but uses simple convolutional blocks instead of residual blocks (Supplementary Fig. 1g). Specifically, each layer is composed of 2 consecutive blocks implementing 3×3 convolutions, each followed by ReLU activations and dropout for network regularization (Supplementary Fig. 1h). The output layer is a single-channel 1×1 convolution block. The stride is equal to 1 in all the convolutional layers and 2 in the max pooling and transposed convolution blocks. All the convolutional kernels were initialized using the He initialization^51^. The size of the filter kernels in the first layer and the dropout rate were considered as hyperparameters and were optimized using Bayesian optimization. The U-Net is analogous to the 3D-U-Net except for the absence of the 3-D convolutional filters at the input. These two networks were independently trained and optimized to separately analyze the effect on the reconstruction performance of the input 3-D convolutional filters and of the residual shortcut connections. In addition, we trained and optimized a network with the same characteristics as the above U-Net to perform the post-processing of power Doppler images that were generated by conventional processing of sparse compound sequences (PP-U-Net).

### Datasets

We trained the networks to learn a function *y = f(x)* that maps the input sequence *x* of compound frames of *N*_*x*_*×N*_*y*_ pixels to the output power Doppler image *y* of dimensions *N*_*x*_*×N*_*y*_ (Fig. 1b). In all our experiments, we used images of 96×96 pixels, and we normalized the input compound datasets. SoA images were obtained from *in-vivo* acquisitions of coronal slices of the rat brain reconstructed by state-of-the-art power Doppler processing using 250 complex compound frames (see “Power Doppler processing” section). To improve the network regularization, we performed random cropping when more than 96 pixels were available in any image dimension, and a random horizontal flipping was applied with a probability of 50%. In total, we used 740 pairs of compound data and power Doppler images for training, 40 pairs for validation, and 40 pairs for testing the reconstruction performance.

We performed under-sampling of the compound sequences in the temporal domain by selecting the first *k* frames in each sequence (Fig. 1b). We retained only the real part of the beamformed data. For the experiments in Fig. 4, we also under-sampled the compound frames in the image domain by selecting sub-samples of pixels with a ratio *m = N*_*Ret*_*/N*_*Tot*_, with *N*_*Ret*_ the number of retained pixels and *N*_*Tot*_ = 96^2^ the number of total image pixels (*m* = 1/2 in Supplementary Fig. 5a and *m* = 1/4 in Supplementary Fig. 5b).

We calculated the compression factor as

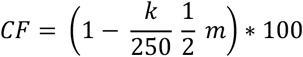

where the factor of 1/2 accounts for the missing imaginary part.

### Training and hyperparameter optimization

At each iteration, the networks predict a new estimate *ŷ*_*i*_, and the parameters are learned using the Adam optimizer^52^ with β_1_ = 0.9, β_2_ = 0.999, and ε = 10^−7^ to minimize the loss function

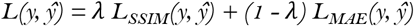

with

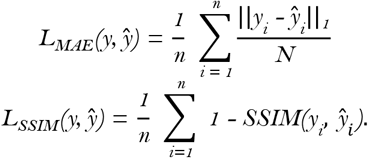

In the above equations, y denotes the SoA images, ‖· ‖_1_ the l_1_ norm, *N* the number of image pixels, and *n* the number of examples. The structural dissimilarity index metric loss L_SSIM_ is a perceptual loss based on the structural similarity index metric (SSIM), which integrates luminance, contrast, and structural information^53,54^. A kernel of 3×3 pixels was used for the SSIM calculation. We considered the learning rate and the parameter λ as hyperparameters, and their optimal value was determined via Bayesian optimization.

We based our quantitative performance analysis on the SSIM of the reconstructed images versus the respective SoA images, the normalized mean squared error NMSE = ‖y_i_ - ŷ_i_‖_2_ / ‖y_i_‖_2_, with ‖· ‖_2_ the *l*_*2*_ norm, and on the peak signal-to-noise ratio (PSNR). We implemented the networks in Python using TensorFlow 2.1 with Keras API. The networks were trained on a single NVIDIA Titan RTX GPU with 24 GB of RAM. The mini-batch size was set to 1 in all the experiments.

For each network, we first optimized the hyperparameters using the Bayesian optimization routine in the Keras Tuner library. We ran 15 optimization trials using the sparse dataset with CF 75%. The optimization routine was instructed to maximize the validation SSIM. Each trial trained the reconstruction CNNs for 2500 epochs and selected the model with the best performance. The results of the optimal hyperparameter search for all the networks are reported in Supplementary Table 1. Then, we trained the CNNs with the optimal hyperparameters using CFs of 80%, 85%, 90%, and 95%. We trained the Res-U-Net and 3D-U-Net networks for 1500 epochs (we noted that these CNNs converged faster during optimization) and the U-Net for 2500 epochs. Five-hundred epochs were used for the optimization and training of the PP-U-Net. In all trainings, the model with the best validation SSIM was saved.

### Ultrasound system and data acquisition

For ultrasound data acquisition, we used two 128-element linear array transducers (L22-14vX and L22-14vLF; Verasonics Inc.) operating at a 15-MHz center frequency with a Vantage 256 research scanner (Verasonics Inc.). The probes are geometrically identical apart from the focus in the elevation plane; the L22-14vX is focused at a distance of 8 mm, and the L22-14vLF is focused at 20 mm. For exact positioning relative to the skull landmarks, the imaging probe was housed in a custom 3-D printed holder mounted on a motorized positioning system. Ultrasound gel was used for acoustic coupling. We used tilted plane waves at angle (−6°, −3°, 0°, 3°, 6°) emitted with a pulse repetition frequency of 19 kHz. Two plane waves were averaged for each angle to increase the signal-to-noise ratio, giving a total of 10 emissions per compound frame. We acquired data for 250 compound frames at a rate of 1 kHz (i.e., a new sequence of compound frames every 250 ms), and the data for each compound sequence (250·10 emissions) were transferred in batch to the host computer. Compound frames were created by beamforming the received sensor RF data in a regular grid of pixels of 100×100 μm^2^ in an NVIDIA Titan RTX GPU using a GPU beamformer^55^. Ultrasound data were acquired asynchronously and continuously, i.e., a new sequence of frames was acquired during processing of the previous sequence and held in the scanner buffer until the host computer was available. The compound frames were saved on the host machine for offline processing. The final power Doppler frame rate was 0.6 frames/sec.

### Conventional power Doppler processing

Sequences of compound ultrasound frames were processed in Matlab (MathWorks) for clutter filtration and power Doppler computation. We used a 5^th^-order temporal high-pass Butterworth filter with a cutoff frequency of 40 Hz cascaded with an SVD filter that eliminates the first singular value^8^. In the Doppler space, frequencies are linearly proportional to the velocity of the scatterers from which the Doppler signal originated. Therefore, it is expected that signals emanating from the slowly moving tissue surrounding the blood vessels (clutter) are positioned at around 0 Hz, and this assumption justifies the use a temporal high-pass filter. Singular value decomposition filters are instead based on the assumption that, while blood signals are highly incoherent due to the time-varying stochastic distribution of the moving scatterers (red blood cells), tissue signals maintain a high degree of correlation over time, and therefore aim to eliminate the highly coherent components. At each pixel location (x, y), the intensity of the filtered signal **s**_Blood_(x, y, t) was then calculated to find the power Doppler value *I(x, y) = ∫* ***s***^*2*^*(x, y, t) dt*. For the SoA processing (250 complex compound frames), the entire time window of 250 ms was integrated.

### Animal preparation and imaging experiments

The experimental protocol for the animal study was approved by the Institutional Animal Care and Use Committee at Stanford University. The university’s animal care and use program and facilities are AAALAC International accredited, PHS-assured, and USDA licensed. Long Evans and Sprague Dawley rats (Charles River; n = 15; age 10-14 weeks; weight 260-400 g) were used in this study. We prepared the animals by performing a bilateral surgical craniotomy and chronic prosthesis implant using previously published protocols^7^. Briefly, animals were anesthetized with 3.5% isoflurane and anesthesia was maintained with 1.5% isoflurane. Rats were placed in a stereotaxic frame during surgery for head fixation and orientation. Body temperature was monitored by a rectal probe and maintained at 36.5 °C using a warming pad (RightTemp Jr.; Kent Scientific). A pulse oximeter was used to monitor heart rate and arterial oxygen saturation (MouseStat Jr.; Kent Scientific). We administered anti-inflammatory to prevent brain swelling and inflammation (1 mg/kg dexamethasone intraperitoneally). After a skin incision was performed, parietal and frontal skull bone fragments (AP +4 to −9 mm; ML ±6 mm) were cut using a handheld high-speed drill with a 0.7 mm drill bit (Fine Science Tools). We gently removed the bone flaps, paying special attention to avoid any damage to the dura mater. We used dental cement (Tetric EvoFlow; Ivoclar Vivadent) to seal a 125 μm thick polymethylpentene prosthesis covering the entire craniotomy. The bone was pre-treated with a bonding agent (iBOND Total Etch; Kulzer). The space between the dura mater and the polymer prosthesis was filled with 0.9% sterile saline. Animals were then allowed to recover for 1 week before the first imaging session.

During the imaging sessions, animals were either anesthetized and kept under anesthesia with 1.5% isoflurane while placed in a stereotaxic frame or were lightly sedated with 0.5% isoflurane and kept in a restraining apparatus^56^. The restrained imaging protocol was also used in the lightly sedated fUS experiment of Fig. 5.

### Visual stimulation experiment

In the visual-task-evoked experiment, rats were anesthetized, placed in a stereotaxic frame, and kept in a dark chamber for at least 30 min prior to the visual stimulation session for dark adaptation. Bilateral visual stimuli were delivered using two green light LEDs driven by a custom power supply circuit. We controlled the stimulus pattern through a microcontroller board (Arduino Uno) connected to Matlab via the serial port and interfaced with the Verasonics scanner for synchronization with the imaging sequence. For each light stimulus, the LEDs were flashed for 30 s at a frequency of 3 Hz. Each stimulus was followed by a >30 s pause in a pseudo-random fashion. This stimulation protocol was shown to maximize visual cortex response in prior fUS imaging studies^57^.

### Functional activation maps

Functional activation maps were created by calculating the Pearson’s correlation coefficient *r* between the temporal power Doppler signal and the stimulus pattern^1,8^. We used a Fisher’s transformation to calculate the *z* score as

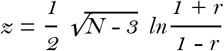

with *N* the number of temporal samples. Each pixel was considered significant for *z* > 3.1, which corresponds to *P* < 0.001 in a one-tailed *t*-test. We used this threshold to create binary activation maps that show only the significant pixels.

## Data availability

The code to reproduce the results of the paper is available at https://github.com/todiian/deep-fus.

## Acknowledgements

This work was supported by a Seed Grant from the Stanford Wu Tsai Neurosciences Institute and the NIH BRAIN Initiative (NIH/NIMH RF1MH114252) and HEAL Initiative (NIH/NINDS UG3NS115637). TDI was supported by a Stanford University School of Medicine Dean’s Postdoctoral Fellowship. We would like to thank Jeremy Dahl, PhD for support with the ultrasound equipment and Dongwoon Hyun, PhD for technical assistance with the GPU beamformer. We would also like to thank Sam Baker, DVM, and Madeline Hughes Cooper for assistance with the surgical protocols. We would like to thank the Airan Lab for helpful discussions.

## Author Contribution

T.D.I. designed and implemented the computational approach for image reconstruction, implemented the functional ultrasound imaging sequences, performed the animal experiments, and developed the relevant scripts for data analysis. R.D.A. provided contextual advising, the hardware for data acquisition, and secured funding support. All authors together drafted and reviewed the figures and manuscript.

## References

1. Macé, E. et al. Functional ultrasound imaging of the brain. Nat. Methods 8, 662–664 (2011).

2. Macé, É. et al. Whole-Brain Functional Ultrasound Imaging Reveals Brain Modules for Visuomotor Integration. Neuron 100, 1241–1251 (2018).

3. Demené, C. et al. Functional ultrasound imaging of brain activity in human newborns. Sci. Transl. Med. 9, (2017).

4. Imbault, M., Chauvet, D., Gennisson, J. L., Capelle, L. & Tanter, M. Intraoperative Functional Ultrasound Imaging of Human Brain Activity. Sci. Rep. 7, 1–7 (2017).

5. Soloukey, S. et al. Functional Ultrasound (fUS) During Awake Brain Surgery: The Clinical Potential of Intra-Operative Functional and Vascular Brain Mapping. Front. Neurosci. 13, 1–14 (2020).

6. Baranger, J. et al. Bedside functional monitoring of the dynamic brain connectivity in human neonates. Nat. Commun. 12, 1080 (2021).

7. Sieu, L.-A. et al. EEG and functional ultrasound imaging in mobile rats. Nat. Methods 12, 831–834 (2015).

8. Urban, A. et al. Real-time imaging of brain activity in freely moving rats using functional ultrasound. Nat. Methods 12, 873–878 (2015).

9. Osmanski, B.-F., Pezet, S., Ricobaraza, A., Lenkei, Z. & Tanter, M. Functional ultrasound imaging of intrinsic connectivity in the living rat brain with high spatiotemporal resolution. Nat. Commun. 5, 5023 (2014).

10. Ferrier, J., Tiran, E., Deffieux, T., Tanter, M. & Lenkei, Z. Functional imaging evidence for task-induced deactivation and disconnection of a major default mode network hub in the mouse brain. Proc. Natl. Acad. Sci. U. S. A. 117, 15270–15280 (2020).

11. Bergel, A., Deffieux, T., Demené, C., Tanter, M. & Cohen, I. Local hippocampal fast gamma rhythms precede brain-wide hyperemic patterns during spontaneous rodent REM sleep. Nat. Commun. 9, 5364 (2018).

12. Blaize, K. et al. Functional ultrasound imaging of deep visual cortex in awake nonhuman primates. Proc. Natl. Acad. Sci. U. S. A. 117, 14453–14463 (2020).

13. Dizeux, A. et al. Functional ultrasound imaging of the brain reveals propagation of task-related brain activity in behaving primates. Nat. Commun. 10, 1–9 (2019).

14. Norman, S. L. et al. Single-trial decoding of movement intentions using functional ultrasound neuroimaging. Neuron 109, 1–13 (2021).

15. Demené, C. et al. Spatiotemporal Clutter Filtering of Ultrafast Ultrasound Data Highly Increases Doppler and fUltrasound Sensitivity. IEEE Trans. Med. Imaging 34, 2271–2285 (2015).

16. Baranger, J. et al. Adaptive Spatiotemporal SVD Clutter Filtering for Ultrafast Doppler Imaging Using Similarity of Spatial Singular Vectors. IEEE Trans. Med. Imaging 37, 1574–1586 (2018).

17. Rabut, C. et al. 4D functional ultrasound imaging of whole-brain activity in rodents. Nat. Methods 16, 994–997 (2019).

18. Sauvage, J. et al. 4D Functional Imaging of the Rat Brain Using a Large Aperture Row-Column Array. IEEE Trans. Med. Imaging 39, 1884–1893 (2020).

19. Ang, E. S. B. C., Gluncic, V., Duque, A., Schafer, M. E. & Rakic, P. Prenatal exposure to ultrasound waves impacts neuronal migration in mice. Proc. Natl. Acad. Sci. U. S. A. 103, 12903–12910 (2006).

20. Ronneberger, O., Fischer, P. & Brox, T. U-net: Convolutional networks for biomedical image segmentation. Lect. Notes Comput. Sci. (including Subser. Lect. Notes Artif. Intell. Lect. Notes Bioinformatics) 9351, 234–241 (2015).

21. Yang, G. et al. DAGAN: Deep De-Aliasing Generative Adversarial Networks for Fast Compressed Sensing MRI Reconstruction. IEEE Trans. Med. Imaging 37, 1310–1321 (2018).

22. Davoudi, N., Deán-Ben, X. L. & Razansky, D. Deep learning optoacoustic tomography with sparse data. Nat. Mach. Intell. 1, 453–460 (2019).

23. Guan, S., Khan, A. A., Sikdar, S. & Chitnis, P. V. Fully Dense UNet for 2-D Sparse Photoacoustic Tomography Artifact Removal. IEEE J. Biomed. Heal. Informatics 24, 568–576 (2020).

24. Jin, K. H., Mccann, M. T., Froustey, E. & Unser, M. Deep Convolutional Neural Network for Inverse Problems in Imaging. IEEE Trans. Image Process. 26, 4509–4522 (2017).

25. Han, Y. & Ye, J. C. Framing U-Net via Deep Convolutional Framelets: Application to Sparse-View CT. IEEE Trans. Med. Imaging 37, 1418–1429 (2018).

26. Luchies, A. C. & Byram, B. C. Deep Neural Networks for Ultrasound Beamforming. IEEE Trans. Med. Imaging 37, 2010–2021 (2018).

27. Hyun, D., Brickson, L. L., Looby, K. T. & Dahl, J. J. Beamforming and speckle reduction using neural networks. IEEE Trans. Ultrason. Ferroelectr. Freq. Control 66, 898–910 (2019).

28. Hyun, D. et al. Nondestructive Detection of Targeted Microbubbles Using Dual-Mode Data and Deep Learning for Real-Time Ultrasound Molecular Imaging. IEEE Trans. Med. Imaging 1–11 (2020). doi:10.1109/TMI.2020.2986762

29. Youn, J. et al. Detection and Localization of Ultrasound Scatterers Using Convolutional Neural Networks. IEEE Trans. Med. Imag. (2020). doi:10.1109/TMI.2020.3006445

30. Gasse, M. et al. High-quality plane wave compounding using convolutional neural networks. IEEE Trans. Ultrason. Ferroelectr. Freq. Control 64, 1637–1639 (2017).

31. Yoon, Y. H., Khan, S., Huh, J. & Ye, J. C. Efficient B-Mode Ultrasound Image Reconstruction From Sub-Sampled RF Data Using Deep Learning. IEEE Trans. Med. Imaging 38, 325–336 (2019).

32. Luijten, B. et al. Adaptive Ultrasound Beamforming using Deep Learning. IEEE Trans. Med. Imag. 1–12 (2019). doi:10.1109/TMI.2020.3008537

33. Nair, A. A., Washington, K. N., Tran, T. D., Reiter, A. & Bell, M. A. L. Deep learning to obtain simultaneous image and segmentation outputs from a single input of raw ultrasound channel data. IEEE Trans. Ultrason. Ferroelectr. Freq. Control (2020). doi:10.1109/tuffc.2020.2993779

34. Paxinos, G. & Watson, C. The Rat Brain in Stereotaxic Coordinates: Hard Cover Edition. (Academic Press, 1998).

35. Zhu, B., Liu, J. Z., Cauley, S. F., Rosen, B. R. & Rosen, M. S. Image reconstruction by domain-transform manifold learning. Nature 555, 487–492 (2018).

36. Vishnevskiy, V., Walheim, J. & Kozerke, S. Deep variational network for rapid 4D flow MRI reconstruction. Nat. Mach. Intell. 2, 228–235 (2020).

37. Shen, L., Zhao, W. & Xing, L. Patient-specific reconstruction of volumetric computed tomography images from a single projection view via deep learning. Nat. Biomed. Eng. 3, 880–888 (2019).

38. Wiacek, A., Gonzalez, E. & Bell, M. A. L. CohereNet: A Deep Learning Architecture for Ultrasound Spatial Correlation Estimation and Coherence-Based Beamforming. IEEE Trans. Ultrason. Ferroelectr. Freq. Control (2020). doi:10.1109/tuffc.2020.2982848

39. Huijben, I. A. M., Veeling, B. S., Janse, K., Mischi, M. & van Sloun, R. J. G. Learning Sub-Sampling and Signal Recovery with Applications in Ultrasound Imaging. IEEE Trans. Med. Imag. 0062, 1–13 (2019).

40. Fishell, A. K. et al. Portable, field-based neuroimaging using high-density diffuse optical tomography. Neuroimage 215, (2020).

41. Wang, J. B. et al. Focused Ultrasound for Noninvasive, Focal Pharmacologic Neurointervention. Front. Neurosci. 14, (2020).

42. Wang, J. B., Aryal, M., Zhong, Q., Vyas, D. B. & Airan, R. D. Noninvasive Ultrasonic Drug Uncaging Maps Whole-Brain Functional Networks. Neuron 100, 728–738 (2018).

43. Di Ianni, T., Hemmsen, M. C., Muntal, P. L., Jorgensen, I. H. H. & Jensen, J. A. System-Level Design of an Integrated Receiver Front End for a Wireless Ultrasound Probe. IEEE Trans. Ultrason. Ferroelectr. Freq. Control 63, (2016).

44. Di Ianni, T. et al. A Vector Flow Imaging Method for Portable Ultrasound Using Synthetic Aperture Sequential Beamforming. IEEE Trans. Ultrason. Ferroelectr. Freq. Control 64, (2017).

45. Rabut, C. et al. Pharmaco-fUS: Quantification of pharmacologically-induced dynamic changes in brain perfusion and connectivity by functional ultrasound imaging in awake mice. Neuroimage 222, 117231 (2020).

46. Ma, Z., Wang, F., Wang, W., Zhong, Y. & Dai, H. Deep learning for in vivo near-infrared imaging. Proc. Natl. Acad. Sci. 118, e2021446118 (2021).

47. O’Reilly, M. A., Muller, A. & Hynynen, K. Ultrasound Insertion Loss of Rat Parietal Bone Appears to Be Proportional to Animal Mass at Submegahertz Frequencies. Ultrasound Med. Biol. 37, 1930–1937 (2011).

48. Errico, C. et al. Transcranial functional ultrasound imaging of the brain using microbubble-enhanced ultrasensitive Doppler. Neuroimage 124, 752–761 (2016).

49. Maresca, D. et al. Acoustic biomolecules enhance hemodynamic functional ultrasound imaging of neural activity. Neuroimage 209, 0–7 (2020).

50. Demené, C. et al. Transcranial ultrafast ultrasound localization microscopy of brain vasculature in patients. Nat. Biomed. Eng. 5, 219–228 (2021).

51. He, K., Zhang, X., Ren, S. & Sun, J. Deep residual learning for image recognition. Proc. IEEE Comput. Soc. Conf. Comput. Vis. Pattern Recognit. 770–778 (2016). doi:10.1109/CVPR.2016.90

52. Kingma, D. P. & Ba, J. L. Adam: A method for stochastic optimization. 3rd Int. Conf. Learn. Represent. ICLR 2015 - Conf. Track Proc. 1–15 (2015).

53. Wang, Z., Bovik, A. C., Sheikh, H. R. & Simoncelli, E. P. Image quality assessment: From error visibility to structural similarity. IEEE Trans. Image Process. 13, 600–612 (2004).

54. Zhao, H., Gallo, O., Frosio, I. & Kautz, J. Loss Functions for Image Restoration With Neural Networks. IEEE Trans. Comput. Imaging 3, 47–57 (2016).

55. Hyun, D., Trahey, G. E. & Dahl, J. J. Real-time high-framerate in vivo cardiac SLSC imaging with a GPU-based beamformer. 2015 IEEE Int. Ultrason. Symp. IUS 2015 1–4 (2015). doi:10.1109/ULTSYM.2015.0077

56. Stenroos, P. et al. Awake rat brain functional magnetic resonance imaging using standard radio frequency coils and a 3D printed restraint kit. Front. Neurosci. 12, (2018).

57. Gesnik, M. et al. 3D functional ultrasound imaging of the cerebral visual system in rodents. Neuroimage 149, 267– 274 (2017).

