## Supplementary Information for "Deep-fUS: Functional ultrasound imaging of the brain using deep learning and sparse data"

### Supplementary Figures

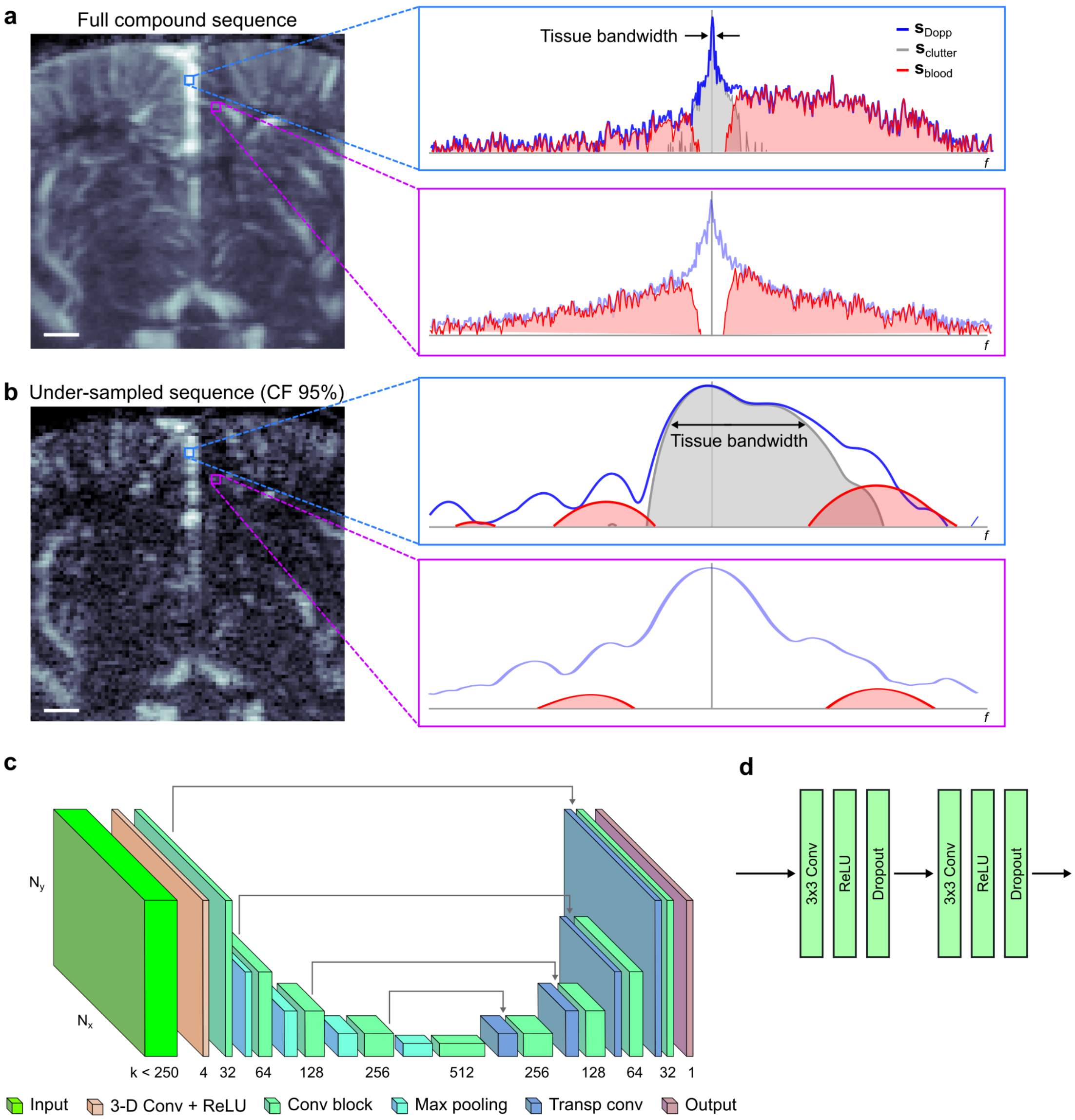

**Supplementary Figure 1**: **a**, Coronal power Doppler image of the rat brain reconstructed with the state-of-the-art approach using 250 complex compound frames and spectra of the Doppler signal in correspondence of a larger blood vessel (Top) and a smaller blood vessel (Bottom). The frequency axis is proportional to the Doppler velocity. The red shaded area is proportional to the power Doppler signal. **b**, Same power Doppler image reconstructed using an under-sampled sequence with 95% compression factor (CF) and spectra of the Doppler signal in the same locations as in **a**. With shorter acquisition sequences, tissue clutter filtration is especially challenging in minor vessels due to the broadening of the tissue bandwidth and the overlapping of the tissue and blood spectra. **c**, Schematic representation of the 3D-U-Net model. **d**, Convolutional blocks are composed of two cascaded Conv/ReLU/Dropout layers. Scale bar in **a**, **b**: 1 mm.

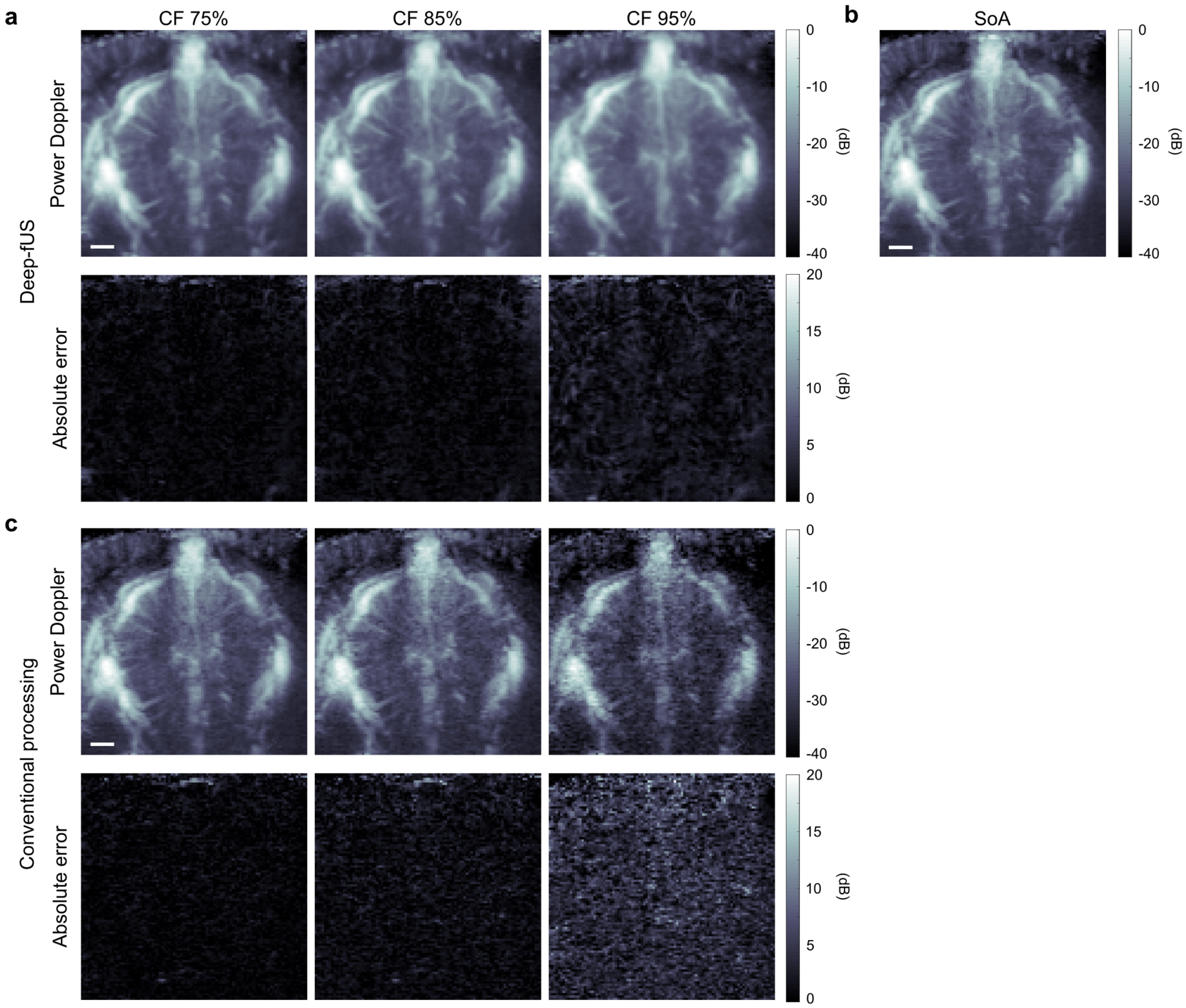

**Supplementary Figure 2**: **a**, Representative power Doppler image of a coronal slice of the rat brain reconstructed by Deep-fUS with compression factor (CF) between 75% and 95% (Top) and absolute error images calculated versus the state-of-the-art (SoA) image (Bottom). **b**, SoA image reconstructed with the conventional processing using 250 complex compound frames. **c**, Power Doppler images reconstructed with the conventional approach from under-sampled compound data (Top) and respective absolute error images (Bottom). Scale bar: 1 mm.

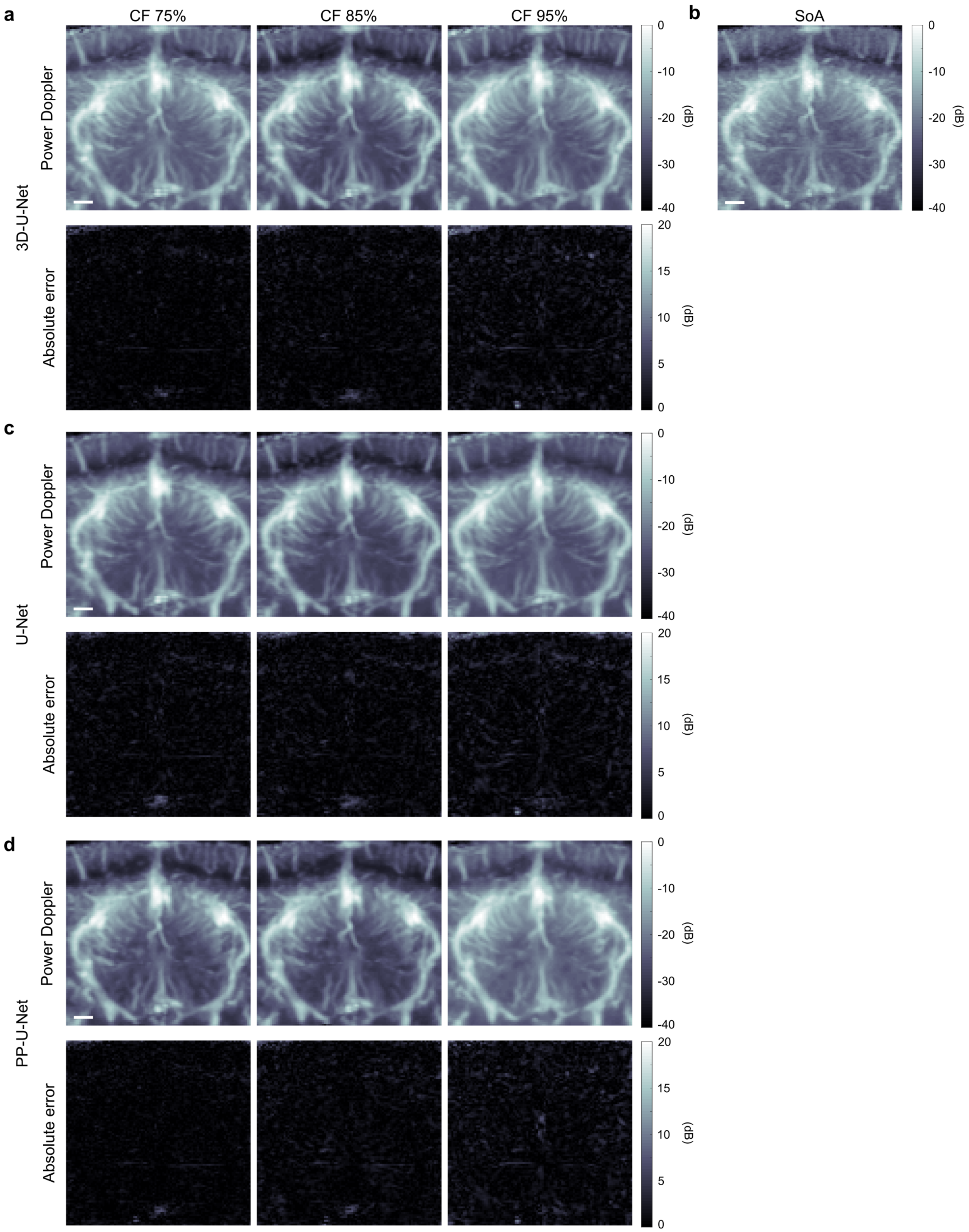
a
CF 75%
CF 85%
CF 95%
b
SoA
3D-U-Net
Power Doppler
Absolute error
c
U-Net
d
PP-U-Net
(dB)
0
-10
-20
-30
-40
20
15
10
5
0

**Supplementary Figure 3**: Coronal slice of the rat brain reconstructed by the (**a**) 3D-U-Net, (**c**) U-Net, and (**d**) PP-U-Net networks with compression factor (CF) between 75% and 95% (Top) and absolute error images calculated versus the state-of-the-art (SoA) reference (Bottom). **b**, SoA image reconstructed by conventional processing using 250 complex compound frames. Scale bar: 1 mm.

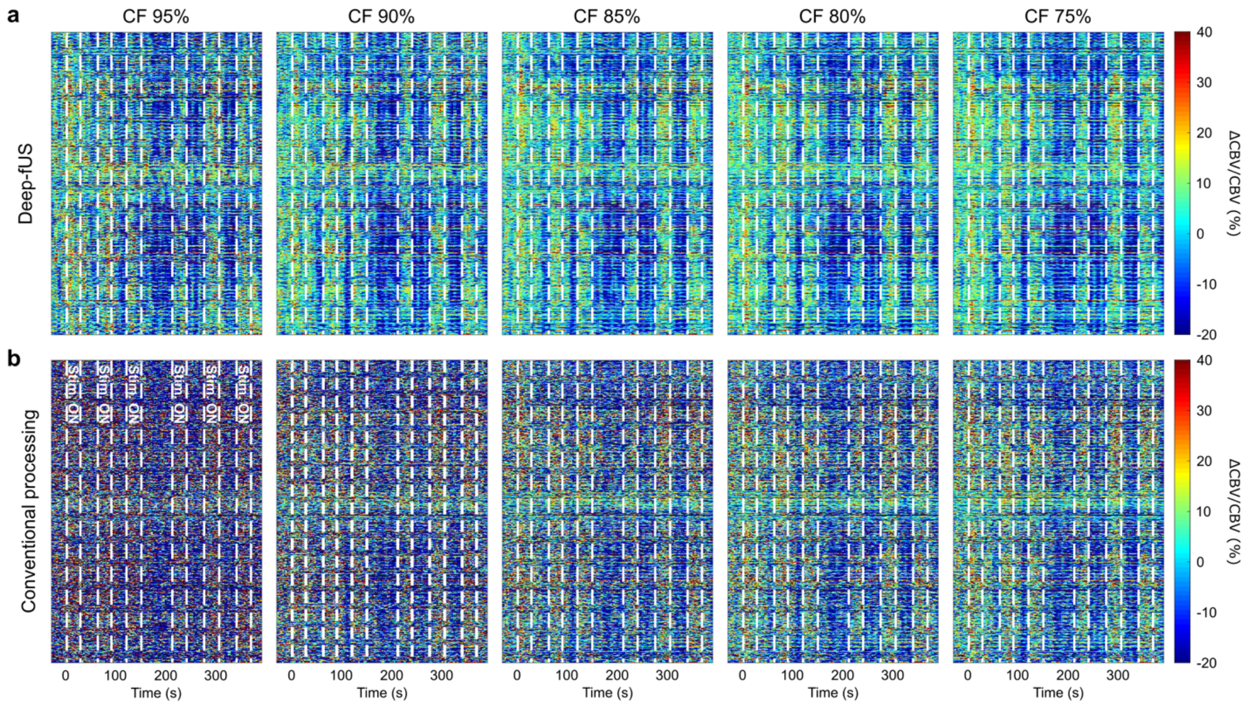

**Supplementary Figure 4**: Relative cerebral blood volume (CBV) temporal signals in all the statistically significant pixels of the SoA map in Fig. 3b for the series reconstructed by Deep-fUS (**a**) and by conventional processing (**b**) with compression factor (CF) between 75% and 95%. The white dashed lines display the ON/OFF stimulus times.

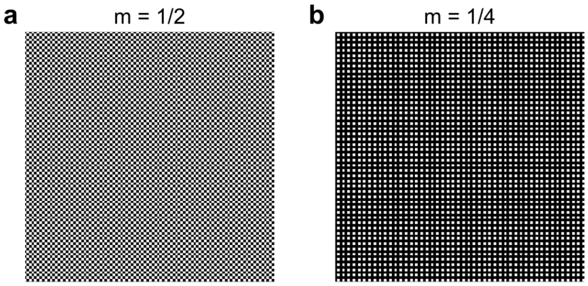

**Supplementary Figure 5**: Sampling maps used to create spatially under-sampled compound frames. The black and white pixels show discarded and retained pixels, respectively. The spatial sampling ratio is m = 1/2 (**a**) and m = 1/4 (**b**).

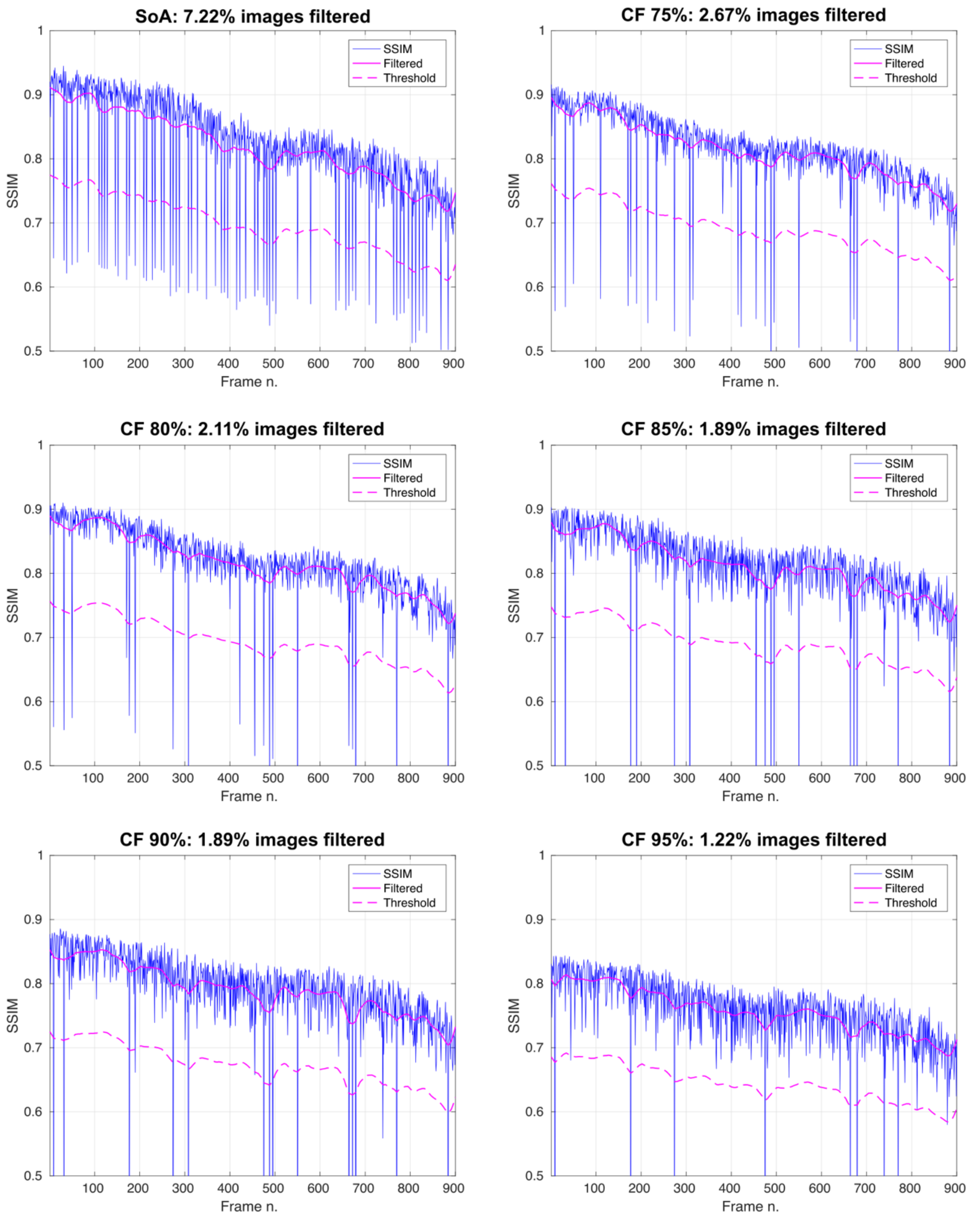

**Supplementary Figure 6**: Structural similarity index metric (SSIM) for each image in the series of power Doppler images from the lightly sedated fUS experiment. SSIM was calculated versus the baseline image (blue) and low-pass filtered to remove the noise (solid magenta). The motion filter threshold was defined as 85% of the SSIM value (dashed magenta). This threshold was used to discard images that were significantly degraded relative to the baseline.

### Supplementary Tables

| Network | Hyperparameter | Optimal value |
|---|---|---|
| Res-U-Net | Conv3D, $k_{1,2}$ | 3 |
| | Conv3D, $k_3$ | 16 |
| | Learning rate | $5.5 \times 10^{-4}$ |
| | Dropout rate | 0.2 |
| | Lambda | 0.1 |
| 3D-U-Net | Conv3D, $k_{1,2}$ | 1 |
| | Conv3D, $k_3$ | 16 |
| | Learning rate | $1.1 \times 10^{-4}$ |
| | Dropout rate | 0.1 |
| | Lambda | 0.9 |
| U-Net | Learning rate | $7.4 \times 10^{-5}$ |
| | Dropout rate | 0.2 |
| | Lambda | 0.8 |
| PP-U-Net | Learning rate | $7.4 \times 10^{-4}$ |
| | Dropout rate | 0.1 |
| | Lambda | 0.2 |

**Supplementary Table 1:** Results of Bayesian hyperparameter optimization.

| Network | | Res-U-Net | 3D-U-Net | U-Net | PP-U-Net |
|---|---|---|---|---|---|
| N. layers | | 5+1 | 5+1 | 5 | 5 |
| N. trainable parameters | | 9,788,421 | 8,773,701 | 8,665,633 | 8,629,921 |
| Training epochs | | 1500 | 1500 | 2500 | 500 |
| Processing time (ms/image) | CF 75% | 13.5 | 7.12 | 5.28 | 2.06 + 255.7 |
| | CF 80% | 11.1 | 5.67 | 4.6 | 2.04 + 220.3 |
| | CF 85% | 9.2 | 4.77 | 3.88 | 2.09 + 207.2 |
| | CF 90% | 6.7 | 3.89 | 3.34 | 2.08 + 194.8 |
| | CF 95% | 4.4 | 2.95 | 2.66 | 2.04 + 183.4 |

**Supplementary Table 2:** Network parameters and processing times for the different models. For the U-Net-PP model, the additional processing time overhead is a result of the pre-processing of under-sampled images. This was computed on a CPU power Doppler implementation. All the processing times were calculated as the average over 390 power Doppler images from the visual task-evoked activation experiment.

| Network | CF | SSIM | PSNR (dB) | NMSE |
|---|---|---|---|---|
| Res-U-Net (recon.) | 75% | 0.9166 ± 0.0174 | 30.2851 ± 1.5516 | 0.0399 ± 0.0222 |
| | 80% | 0.9041 ± 0.0192 | 29.4798 ± 1.5038 | 0.0448 ± 0.0278 |
| | 85% | 0.8915 ± 0.0211 | 28.8217 ± 1.4282 | 0.0506 ± 0.0224 |
| | 90% | 0.8619 ± 0.0264 | 27.8481 ± 1.4058 | 0.0701 ± 0.0472 |
| | 95% | 0.8154 ± 0.0353 | 26.7270 ± 1.2469 | 0.1106 ± 0.0364 |
| 3D-U-Net (recon.) | 75% | 0.9046 ± 0.0189 | 29.5322 ± 1.4163 | 0.0647 ± 0.0259 |
| | 80% | 0.8981 ± 0.0201 | 29.4254 ± 1.2746 | 0.0652 ± 0.0335 |
| | 85% | 0.8874 ± 0.0202 | 29.0893 ± 1.2615 | 0.0728 ± 0.0308 |
| | 90% | 0.86 ± 0.0238 | 28.2881 ± 1.2772 | 0.0913 ± 0.0464 |
| | 95% | 0.8139 ± 0.0373 | 27.1319 ± 0.7524 | 0.1177 ± 0.0495 |
| U-Net (recon.) | 75% | 0.8394 ± 0.0342 | 27.6412 ± 0.7867 | 0.1222 ± 0.0488 |
| | 80% | 0.8348 ± 0.0323 | 27.2672 ± 0.7142 | 0.1375 ± 0.0456 |
| | 85% | 0.8384 ± 0.0325 | 27.3371 ± 0.8499 | 0.1399 ± 0.0385 |
| | 90% | 0.8237 ± 0.0337 | 26.7846 ± 0.8862 | 0.1398 ± 0.0467 |
| | 95% | 0.7927 ± 0.0444 | 26.1352 ± 0.8169 | 0.1946 ± 0.0550 |
| PP-U-Net (post-process.) | 75% | 0.9269 ± 0.0153 | 31.1037 ± 1.2606 | 0.0359 ± 0.0239 |
| | 80% | 0.9165 ± 0.0179 | 30.6757 ± 1.4619 | 0.0396 ± 0.0224 |
| | 85% | 0.902 ± 0.0212 | 30.2345 ± 1.1474 | 0.0459 ± 0.0242 |
| | 90% | 0.876 ± 0.0233 | 29.1286 ± 1.3275 | 0.061 ± 0.0323 |
| | 95% | 0.8226 ± 0.0376 | 27.3628 ± 0.9142 | 0.102 ± 0.0537 |

**Supplementary Table 3:** Quantitative performance metrics calculated versus the respective state-of-the-art images for the different models with increasing compression factor (CF). SSIM: Structural similarity index metric. PSNR: peak signal-to-noise ratio. NMSE: Normalized mean squared error. Quantities are reported as mean ± standard deviation calculated over the test set.

| m | 1/2 | 1/4 | 1 |
|---|---|---|---|
| **k** | 50 | 100 | 25 |
| **CF** | 95% | 95% | 95% |
| **SSIM** | 0.8368 ± 0.0311 | 0.8549 ± 0.0257 | 0.8154 ± 0.0353 |
| **PSNR (dB)** | 27.1588 ± 1.3782 | 28.1827 ± 1.1884 | 26.7270 ± 1.2469 |
| **NMSE** | 0.0782 ± 0.0459 | 0.0606 ± 0.0272 | 0.1106 ± 0.0364 |
| **Activation map MAE** | 0.1425 | 0.1216 | 0.1805 |

**Supplementary Table 4:** Quantitative performance metrics calculated versus the respective state-of-the-art (SoA) images for the experiment with spatially under-sampled compound data. m: spatial under-sampling ratio. k: number of compound images in the sparse sequence. CF: compression factor. SSIM: Structural similarity index metric. PSNR: peak signal-to-noise ratio. NMSE: Normalized mean squared error. Activation map MAE: mean absolute error between the statistically significant correlations in the SoA map and the under-sampled maps in the visual-task-evoked activation experiment. Quantitative performance metrics are reported as mean ± standard deviation calculated over the test set.

### Supplementary Videos

**Supplementary Videos 1-3:** Power Doppler image series reconstructed by Deep-fUS and conventional processing with compression factor (CF) of 75% (Video 1), 85% (Video 2), and 95% (Video 3). State-of-the-art (SoA) reference images (left) are displayed side-by-side with images reconstructed from sparse compound sequences (center and right).

**Supplementary Video 4:** Power Doppler image series and relative CBV variation (superimposed heatmap) for the state-of-the-art (SoA) reference (left), Deep-fUS reconstruction using spatially and temporally under-sampled compound sequences with compression factor (CF) of 95% ($m$ = 1/4, $k$ = 100; center), and conventional processing with CF of 95% (right). Images from the visual-evoked activation session were used to create this video. Binary masks of CBV variation were created by selecting only the pixels statistically significantly correlated with the stimulus pattern. Reproduced at 7.5x.

**Supplementary Video 5:** Power Doppler image series from the fUS imaging experiment in a lightly sedated rat. Images were reconstructed using the state-of-the-art approach (SoA; left) and by Deep-fUS with compression factors (CF) of 75% (center) and 95% (right). Images that were discarded by the motion filter are shown with a white border. Using shorter compound sequences reduces the occurrence of motion artifacts and decreases image scrubbing.

### Supplementary Note

#### Calculation of RF and beamformed data throughputs

Considering compound frames composed of 10 plane wave emissions acquired with an array of 128 elements, sampled with a sampling frequency of 60 MHz (4× the pulse center frequency of 15 MHz), and covering a depth of 9.6 mm, the data throughput for each compound frame is

$$\frac{2 *\ 9.6\ 10^{-3}}{1.54\ 10^{3}} * 60\ 10^{6} * 128 * 10 = \ 957{,}510$$

RF samples. This equation considers a speed of sound of 1540 m $s^{-1}$, and the factor 2 accounts for the pulse-echo time-of-flight. Assuming that 250 compound frames are used to compute a power Doppler image, the total number of RF samples to be transmitted from the scanner to the host computer is

$$957{,}510 * 250 = 239{,}377{,}500$$

per image. If 96×96 pixels are beamformed in each compound frame, the beamforming load is

$$96 * 96 * 2 * 250 = 4{,}608{,}000$$

pixels per power Doppler image. The factor of 2 in the last equation accounts for the beamforming of complex (real and imaginary) samples.

By saving beamforming data instead of RF samples, memory usage can be reduced by

$$\frac{957{,}510}{96 * 96 * 2} = 52$$

times, assuming that data are saved with the same precision.

Our CNN processing approach reduces the hardware requirements for data acquisition, as well as the storage and beamforming requirements. For example, in the case of 80% compression factor with $k$ = 100 real compound frames per power Doppler image and no spatial under-sampling ($m$ = 1), the number of RF samples is reduced to

$$957{,}510 * 100 = 95{,}751{,}000$$

(60% reduction), while the number of beamformed pixels to be processed and stored in memory is reduced to

$$96 * 96 * 100 = 921{,}600$$

(80% reduction). With $k$ = 100 and $m$ = 1/4 (Fig. 4), the number of RF samples is unchanged while the number of beamformed pixels becomes

$$\frac{1}{4} * 96 * 96 * 100 = 230{,}400$$

(95% reduction). For the sake of simplicity, in the main text we only refer to the beamformed data reduction to compute the data compression factor.
